# Mitotic bookmarking redundancy by nuclear receptors mediates robust post-mitotic reactivation of the pluripotency network

**DOI:** 10.1101/2022.11.28.518105

**Authors:** Almira Chervova, Amandine Molliex, H. Irem Baymaz, Thaleia Papadopoulou, Florian Mueller, Eslande Hercul, David Fournier, Agnès Dubois, Nicolas Gaiani, Petra Beli, Nicola Festuccia, Pablo Navarro

## Abstract

Mitotic bookmarking transcription factors (TFs) are thought to mediate rapid and accurate post-mitotic gene reactivation. However, the loss of individual bookmarking TFs often leads to the deregulation of only a small proportion of their mitotic targets, raising doubts on the significance and importance of their bookmarking function. Here, we used targeted proteomics of the mitotic bookmarking TF ESRRB, an orphan nuclear receptor, to discover an unexpected redundancy among members of the protein superfamily of nuclear receptors. Focusing on the nuclear receptor NR5A2, which together with ESRRB is essential for mouse pluripotency, we demonstrate conjoint bookmarking activity of both factors on promoters and enhancers of a large fraction of active genes, particularly the most rapidly and strongly reactivated ones. Upon fast and simultaneous degradation of both factors during mitotic exit, hundreds of mitotic targets of ESRRB/NR5A2, including key players of the pluripotency network, display attenuated transcriptional reactivation. We propose that redundancy in mitotic bookmarking TFs, especially by nuclear receptors, confers robustness to the reestablishment of gene regulatory networks after mitosis.

## Introduction

During mitosis many transcription factors (TFs) are inactivated or evicted from the chromatin; however, some remain active and bind to their targets^**1**^. While it is thought that this phenomenon, known as mitotic bookmarking, enables daughter cells to promptly resume transcription after mitosis^**2**^, its exact role and significance is unclear. Indeed, only a small subset of bookmarked genes displays a clear, albeit partial, dependence on their respective bookmarking factor for proper reactivation^**3-13**^. This has led to the hypothesis that the effects of mitotic bookmarking TFs might be insignificant and represent a mere consequence of other chromatin properties^**13**^. However, an alternative hypothesis is that of mitotic bookmarking redundancy: distinct TFs may independently bookmark key genes important for cell identity such that the loss of one would be largely inconsequential.

The nuclear receptor ESRRB, a mitotic bookmarking TF in mouse Embryonic Stem (ES) cells^**5**,**10**^, represents an ideal candidate to assess the notion of mitotic bookmarking redundancy. Indeed, while ESRRB binds a third of its interphase targets (around 10,000 regulatory regions)^**5**,**10**^, only 150 genes require ESRRB to be properly reactivated immediately after mitosis^**5**^. Moreover, while ESRRB knock-out ES cells are viable, the simultaneous loss of ESRRB and another nuclear receptor, NR5A2, is incompatible with the maintenance of pluripotency^**14**^. This strong complementarity between ESRRB and NR5A2 showcases the importance of two functionally redundant nuclear receptors. In fact, a high level of redundancy among TFs of the superfamily of nuclear receptors might be expected since they are evolutionarily and structurally related and share highly similar DNA binding motifs^**15-18**^. Moreover, several nuclear receptors have been shown to coat mitotic chromosomes^**19-25**^, even though their engagement in site-specific interactions genome-wide has only been analysed for ESRRB^**5**,**10**^. Together, these observations suggest that cohorts of nuclear receptors could be involved in mitotic bookmarking processes to ensure the proper post-mitotic reactivation of the pluripotency network.

To investigate mitotic bookmarking redundancy from the perspective of ESRRB, we first established that the most recurrent and prevalent proteins with which it associates on mitotic chromatin are nuclear receptors. Second, focusing on NR5A2 we showed that its retention on mitotic chromatin is long-lived and characterised by site-specific interactions at gene regulatory elements harbouring a specific variant of the DNA binding motif of nuclear receptors. Third, we assessed the functional consequence of dual ESRRB/NR5A2 bookmarking using Auxin-inducible degrons: we found both factors to be conjunctly required for efficiently jumpstarting around 1000 genes after mitosis. These ESRRB/NR5A2-responsive genes during the M-G1 transition are collectively downregulated during differentiation, transiently induced in pluripotent compartments of early embryos and enriched for pluripotency regulators. We conclude that nuclear receptors execute the key task of rapidly reinstating after mitosis the gene regulatory networks supporting cell identity.

## Results

### Mitotic association of ESRRB with nuclear receptors

Previous work showed that ESRRB interacts with a large number of chromatin-associated proteins, including chromatin remodellers, members of the basal transcriptional apparatus, pluripotency TFs and other nuclear receptors^**26**,**27**^. We therefore aimed at identifying which proteins are associated with ESRRB in interphase and in mitosis. To do so, we applied ChIP-MS, a technique similar to ChIP-seq but which uses mass spectrometry to identify the factors crosslinked to the immunoprecipitated protein^**28**^: in our case ESRRB **(Fig.S1A)**. Since we previously showed that fixation with disuccinimidyl glutarate (DSG) and formaldehyde significantly improves the detection of TF localisation to mitotic chromosomes^**10**^, we performed three replicate assays in such conditions together with negative IP controls (**Table S1**, see Methods for details). However, we have also shown that formaldehyde alone is enough to detect substantial numbers of ESRRB mitotic binding sites. Therefore, we also performed ChIP-MS after either DSG and formaldehyde or only formaldehyde crosslinking and compared the results to the respective inputs. In agreement with previous reports^**26**,**27**^, we found other pluripotency TFs and members of the Mediator, NuRD and Swi/Snf complexes associated with ESRRB in asynchronous cells **(Fig.1A, Fig.S1B and Table S1)**. All these proteins were, however, largely undetectable in mitosis. Indeed, a more comprehensive analysis **(Fig.S1C)** identified 105 proteins in asynchronous cells, belonging to different functional groups **(Fig.1B, Table S2)**. Of those proteins, we found only 16 that were identified in mitosis, regardless of the crosslinking strategy and strongly enriched as compared to both the control IP and corresponding input (in red in **Fig.1B, Fig.S1C** and **Table S2**). We also identified 21 additional proteins that were enriched as compared to the control IP but not to the input, raising the possibility that these are highly expressed proteins in mitotic cells leading to some level of unspecific detection (in pink in **Fig.1B, Fig.S1C** and **Table S2**). Notably, none of the pluripotency TFs were identified in mitosis; in fact, the vast majority of proteins was undetectable in mitosis for every functional group except for one, nuclear receptors and related factors, which were almost all found associated with ESRRB in both asynchronous and mitotic cells (**Fig.1A** and red and pink coloured proteins in **Fig.1B**). While the mitotic detection of selected proteins in most functional groups is interesting, as it suggests a priming mechanism mediated by ESRRB to recruit other partners after mitosis, the high occurrence of nuclear receptors in mitotic cells is compelling, representing half of the most confident mitotic hits **(Fig.1B)**. The identified nuclear receptors, with roles in ES cells^**29**^ are: ESRRB itself but also the highly related ESRRA; NR0B1 (also known as DAX1), a nuclear receptor lacking a DNA-binding domain; NR5A2 (also known as LRH1), whose functional role together with ESRRB has been already demonstrated in ES cells; RXRB, a retinoic acid receptor with established roles in pluripotency and differentiation. In addition, other factors that were reported as interacting with nuclear receptors are also associated with ESRRB in asynchronous and mitotic chromatin, such as NSD1^**30**^, TRIM24^**31**^ and SNW1^**32**^. Among this small set of proteins, and given its major role in ES cells^**14**,**27**^, we focused on NR5A2 for further analyses.

**Fig. 1.**
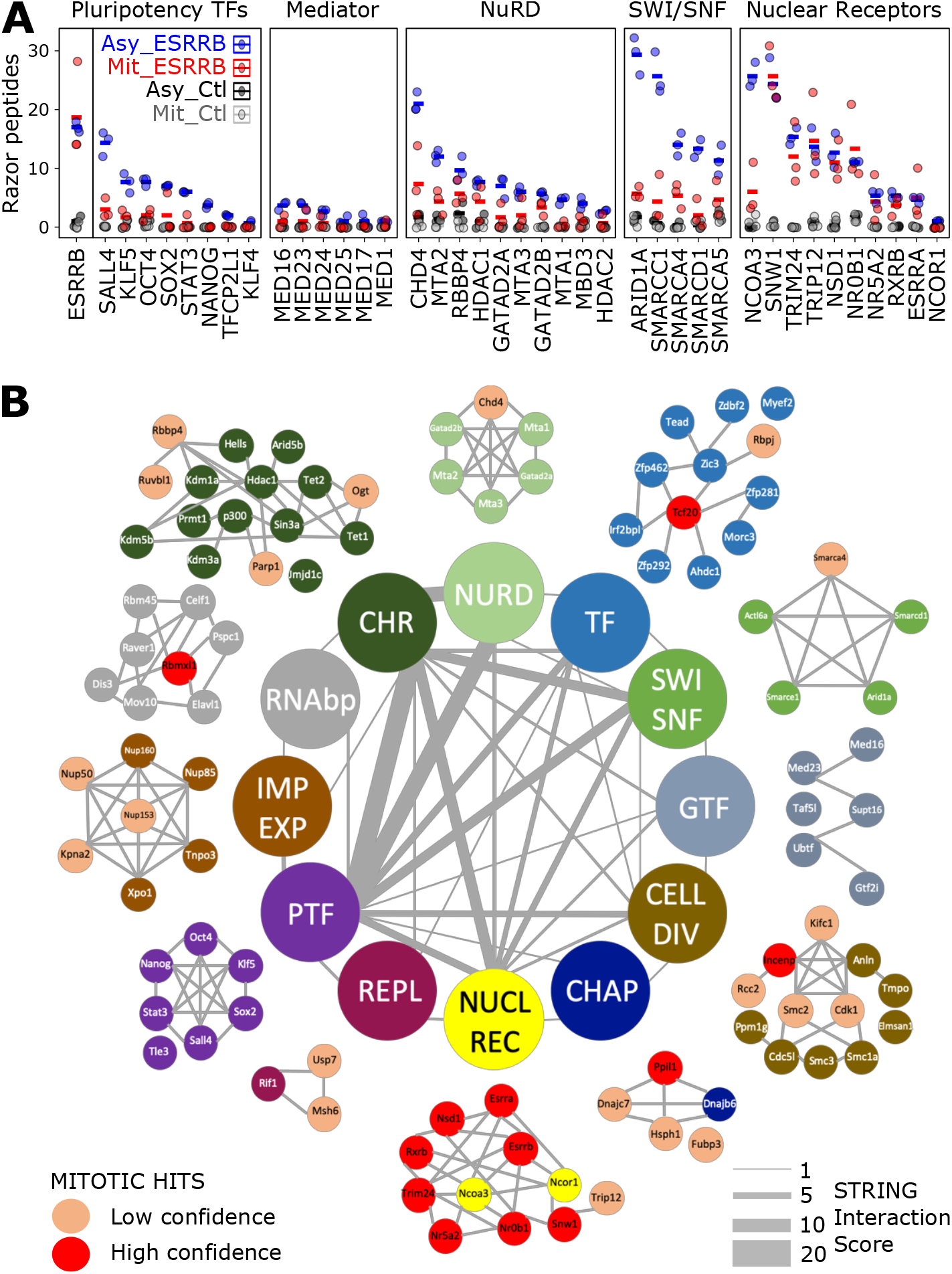
Nuclear receptors are maintained on mitotic chromatin with ESRRB. **(A)** Quantification of peptide abundance for specific examples belonging to different functional groups known to interact with ESRRB in interphase. The number of identified razor peptides is shown; similar observations were made with iBAQ and LFQ quantifications (Tables S1 and S2). **(B)** The ESRRB-centred, chromatin-associated proteome in interphase and mitotic cells, organised by functional groups (NURD: Nucleo-some Remodeling and Deacetylase Complex; TF: Transcription Factors; SWI/SNF: SWItch/Sucrose Non-Fermentable nucleosome remodelling complex; GTF: General Transcription Factors; CELL DIV: proteins with known roles during mitosis; CHAP: proteins with roles in chaperoning activity; NUCL REC: NUCLear RECeptors and known associated factors; REPL: REPLication machinery; PTF: Pluripotency Transcription Factors; IMP EXP: proteins with roles in nuclear IMPort and EXPort; RNAbp: RNA binding proteins; CHR: proteins with known roles in CHRo-matin regulation.

### Long-lived chromatin retention of NR5A2 in mitosis

TFs acting as mitotic bookmarking factors, such as ESRRB, often coat mitotic chromosomes^**5**^, as reproduced here using an ESRRB-TdTomato fusion protein **(Fig.2A)**. As expected from our ChIP-MS results **(Fig.1)**, NR5A2-GFP fusion proteins colocalised with ESRRB on mitotic chromosomes **(Fig.2A)**. Moreover, the mitotic retention of these two factors is mutually independent, as established using previously described KO cell lines^**14**^: in ESRRB KO mitotic cells, NR5A2 remains associated with the chromatin; conversely, in NR5A2 KO cells ESRRB still coats the mitotic chromosomes **(Fig.2B)**. These results are in accord with the large binding independence of the two factors to their common sites on DNA^**14**^. Despite this seemingly similar behaviour, ESRRB and NR5A2 coating of the mitotic chromosomes also displays notable differences: while the chromosomal coating by ESRRB is disrupted by formaldehyde crosslinking^**10**^, as shown for many other TFs^**33**^, that of NR5A2 is retained **(Fig.S2A)**. A potential explanation for this behaviour, already reported for CTCF^**11**^, could be that the residence time of NR5A2 on the chromatin is long, allowing formaldehyde crosslinking^**33**,**10**^. Accordingly, FRAP analyses performed in parallel for ESRRB and NR5A2 clearly show that while fluorescence is rapidly recovered on bleached chromatin for ESRRB, in the case of NR5A2 the recovery is incomplete over 10 seconds **(Fig.2C)**. Even after long times following photobleaching (up to 100s), NR5A2 signal does not recover entirely, indicating that the association of this TF with the chromatin is long-lived. This is particularly evident in interphase, but holds true also in mitosis, where a seemingly immobile fraction of around 25% characterises NR5A2 coating of the mitotic chromosomes **(Fig.2D)**. These results, in sharp contrast to most if not all other bookmarking TFs analysed and more specifically to ESRRB **(Fig.S2B)**, set apart NR5A2 as a TF that is stably associated with the chromatin in interphase and in mitosis. Intrigued by this finding, we asked whether the DNA Binding Domain (DBD) of NR5A2 was sufficient for such long-lived interactions by ectopically expressing DBD-GFP fusion proteins. The recovery of fluorescence for the NR5A2 DBD alone was faster in interphase **(Fig.S2C)**, matching the dynamics of the full protein in mitosis. Addition of the C-terminal domain that contains the Ligand Binding Domain (LBD), responsible for the interaction of nuclear receptors with coactivators, considerably increased the time of fluorescent recovery **(Fig.S2C)**. In mitosis, we found that both the DBD alone and the DBD-LBD fusion proteins efficiently coated the chromosomes **(Fig.S2D)**. Moreover, and in contrast to interphase, the dynamics of fluorescence recovery were found to be similar between the two constructs although the DBD alone displayed increased stability **(Fig.S2C)**. Therefore, we conclude that the DBD alone is sufficient to trigger relatively-long lived chromatin interactions in mitosis, whereas in interphase the LBD, possibly through the mediation of protein-protein interactions, confers further stability to NR5A2 binding to the chromatin.

**Fig. 2.**
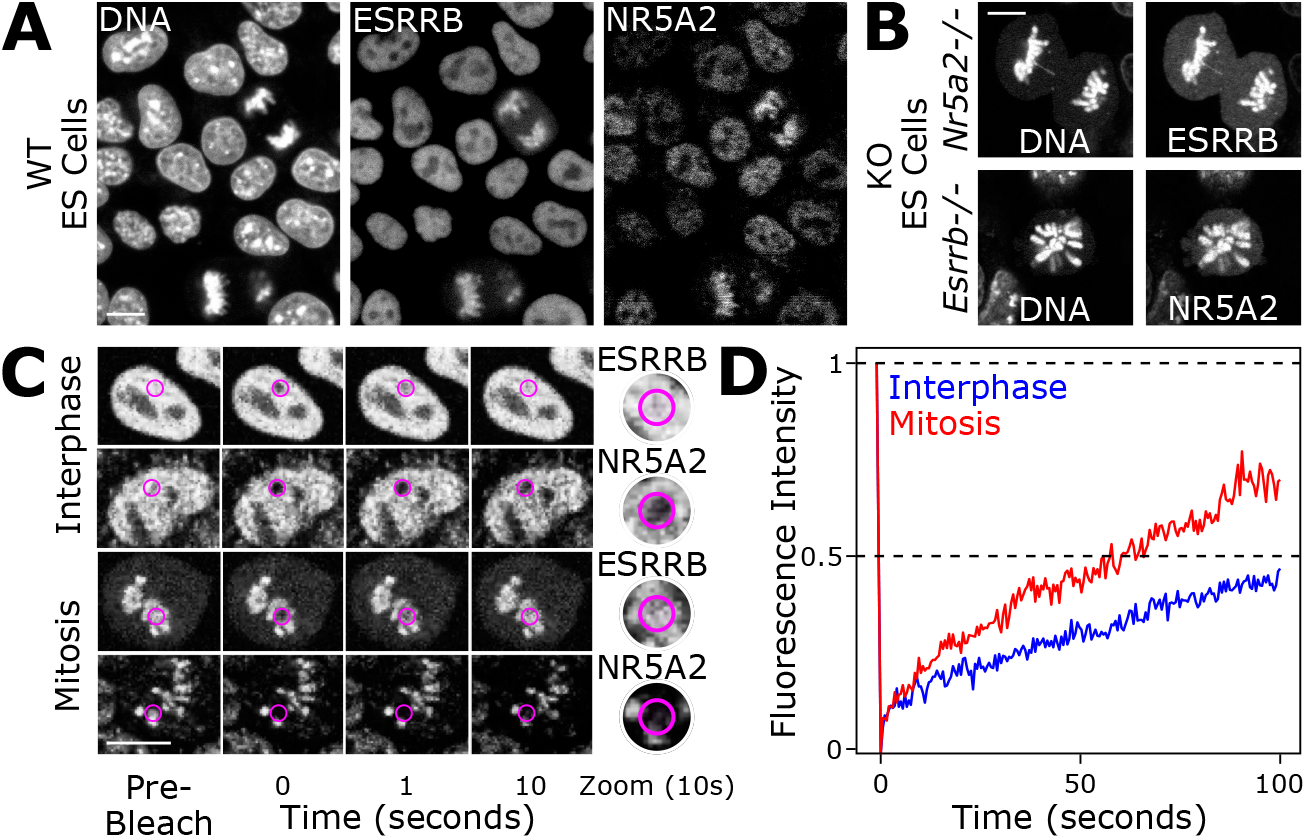
NR5A2 is robustly bound to the chromatin in interphase and during mitosis. **(A)** Live imaging of ESRRB-TdTomato and NR5A2-GFP fusion proteins in wild-type ES cells. Note the mitotic retention of both TFs. **(B)** Live imaging of ESRRB-GFP in NR5A2 KO cells and of NR5A2-GFP in ESRRB KO cells illustrating the mutually independent association with mitotic chromosomes. **(C)** Representative example of FRAP assays in interphase and in mitosis. **(D)** Quantification of the recovery of fluorescence during 100 s following photo-bleaching.

### Mitotic bookmarking by NR5A2

We next aimed at assessing whether NR5A2 engages in site-specific interactions with its DNA targets, a defining property of mitotic bookmarking factors^**1**^. To do this, we performed ChIP-seq assays in asynchronous and highly pure populations of mitotic ES cells. Exploration of the binding profiles throughout the genome confirmed the capacity of NR5A2 to bind mitotic chromatin at specific sites **(Fig.3A)**, which can be described with four main binding trends **(Fig.3B and Table S3)**: regions that are bound by both factors in interphase and in mitosis, thereafter dB (for double bookmarked), regions that are bookmarked by either ESRRB (eB) or NR5A2 (nB) and regions that are bound by the two factors exclusively in asynchronous cells (thereafter lost regions, L). Globally, ESRRB and NR5A2 binding levels were higher at bookmarked regions, including in asynchronous cells **(Fig.S3A,B)**, in keeping with the idea that mitotic bookmarking often takes place at regions of robust binding. The behaviour of ESRRB and NR5A2 is not, however, fully symmetric. Indeed, if at nB regions ESRRB is bound in interphase but mostly lost in mitosis, at eB regions NR5A2 is not found in mitosis simply because this TF is not efficiently recruited at these loci even in interphase **(Fig.3B and Fig.S3A,B)**. This observation, together with our previous finding that NR5A2 and ESRRB show a preference for slightly different DNA motifs^**14**^, prompted us to determine whether distinct sequences are enriched over these 4 groups of regions. For all groups, we found an overrepresentation of the TCAAGGTCA sequence characteristic of Estrogen Related Receptors **(Fig.3C)**, which contains the classical AG-GTCA box typical of many nuclear receptors^**34**,**35**^, extended by a half-site. However, at regions bound by NR5A2 in mitosis (dB and nB), thymidine (T) was less prominent at the seventh base, and as frequent there as cytosine (C). The occurrence of T or C at the seventh position in the motif was already identified as favouring ESRRB versus NR5A2 binding in asynchronous ES cells^**14**^. Analysis of the presence of the motif and more particularly of T/C variants across all regions showed that the efficiency of mitotic bookmarking correlates to the presence of this consensus, with loci containing both T and C variants, or exclusively C, being strongly enriched at dB and nB regions **(Fig.3D)**; in contrast, eB regions were exclusively associated with motifs containing a T. As expected, L regions showed the lowest occurrence of motifs, which almost always contain a T. Moreover, we observed that the type of motifs present in the regions **(Fig.3E)**, as well as their number per region **(Fig.3F)** and quality **(Fig.3G)**, are quantitatively associated with the binding levels of ESRRB and NR5A2, particularly in mitosis. Hence, the average motif score of the regions clearly differentiates bookmarked versus lost status **(Fig.3H)**. Finally, we noted that the motif identified at dB and nB regions was longer, containing an extra AGT at the 5’ end **(Fig.3C)**. While we had already identified the presence of this longer motif in ESRRB bookmarked regions^**5**^, this new analysis clearly associates it with book-marking by NR5A2, with or without ESRRB. Accordingly, the Jaspar database reports a motif highly similar to our long variant as the NR5A2 motif **(Fig.S3C)**. We conclude that the presence of different versions of the ESRRB/NR5A2 motif, together with their degree of similarity to the consensus and their number of occurrences per region, are direct determinants of the behaviour of ESRRB and NR5A2 in mitosis; in interphase, though, ESRRB and NR5A2 can also be recruited by other TFs, often excluded from mitotic chromatin **(Fig.1)**.

**Fig. 3.**
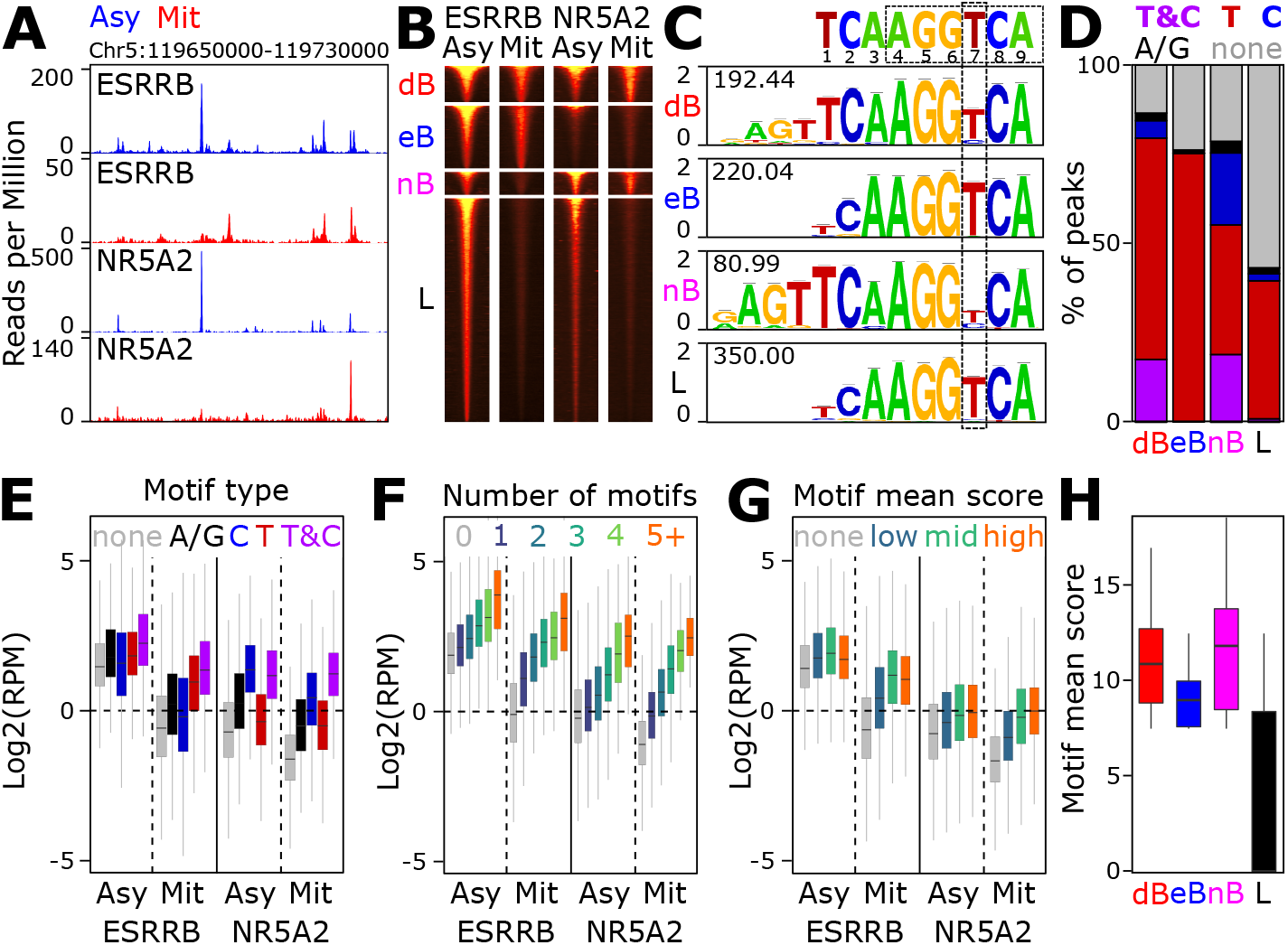
Mitotic bookmarking by NR5A2 of a subset of ESRRB/NR5A2 target loci. **(A)** Representative binding profiles of ESRRB and NR5A2 in asynchronous (Asy) and mitotic (Mit) cells. **(B)** Clustering of ESRRB/NR5A2 binding regions as double bookmarked (dB), bookmarked by ESRRB (eB) or by NR5A2 (nB) only, or exclusive to interphase (L, for lost in mitosis). **(C)** Consensus motifs found de novo at the 4 clusters described in (B), compared to the Orphan Nuclear Receptors cognate sequence. The number within each motif denotes the corresponding -log10(FDR). The 7th base partially discriminating ESRRB (T) versus NR5A2 (C) preferential binding is highlighted. **(D)** Percentage of peaks harbouring both T and C motifs, or exclusively T, C, A/G or no motifs. **(E-G)** Enrichment levels (Reads Per Million) of ESRRB and NR5A2 in asynchronous (Asy) and mitotic (Mit) cells as a function of the presence of different motif types (E), the total number of motifs per region (F) or the mean score of all the motifs per region (G). **(H)** The motif mean score per region shows drastic changes between mitotically bookmarked (dB, eB, nB) and lost (L) regions.

### Chromatin states of ESRRB/NR5A2 binding regions

We next separated ESRRB/NR5A2 binding regions using epigenomic signatures characteristic of active promoters, active enhancers or enhancers lacking marks of activity, and quantified pluripotency TF binding (OCT4, SOX2 and NANOG; **Fig.4A**). At enhancers, we observed that pluripotency TFs were almost exclusively constrained to L regions displaying poor ESRRB/NR5A2 motifs **(Fig.4A)**, indicating that ESRRB and NR5A2 are likely indirectly recruited at these regions. This subset was also characterised by slightly more accessible chromatin and p300 recruitment **(Fig.4A)**. At active enhancers and promoters this trend was not apparent **(Fig.4A)**; instead, we found a small positive correlation between marks of activity and mitotic bookmarking status, with dB globally displaying higher levels of active marks than eB/nB, which in turn showed more enrichment than L regions **(Fig.S4A)**. Moreover, although the effect was rather modest, dB and eB regions also displayed a less pronounced reduction in accessibility in mitosis compared to asynchronous cells **(Fig.S4A)**. We also observed that, proportionally, active promoters display the highest frequency of bookmarked regions (50% against 20-30% for other elements, **Fig.4A** and **Fig.S4B**). This is reflected by a 2 to 3-fold enrichment of promoters within dB and eB regions, which nevertheless remain in absolute terms less frequently bound by these factors than enhancers **(Fig.S4B)**. Overall, this analysis indicates that at enhancers losing ESRRB/NR5A2 in mitosis, NANOG, OCT4 and SOX2 are likely key regulators triggering ESRRB/NR5A2 recruitment in interphase and a modest enrichment for epigenomic features associated with activity. In contrast, at active regulatory elements, ESRRB and NR5A2 have a positive impact in pluripotency TF binding and enrichment for active marks, particularly at regions where ESRRB /NR5A2 engage in mitotic bookmarking. Next, we turned to the analysis of nucleosome organisation **(Fig.4B)**, since we had previously shown that ESRRB and CTCF bookmarked regions maintain nucleosome order in mitosis^**10**,**11**^. In this regard, nB regions are particularly interesting as they were previously considered as lost regions. Using previously published MNase-seq datasets^**10**^, we observed that at all bookmarked regions (dB, eB and nB), the nucleosomes were ordered as nucleosomal arrays in both interphase and mitosis **(Fig.4B)**. Whilst the best phasing of the nucleosomes and the more pronounced nucleosome depleted region over the motif were observed at dB regions, both eB and more significantly nB regions displayed substantial maintenance of nucleosome organisation. In contrast, L regions showed poor organisation and a clear central accumulation of nucleosomes, specifically in mitosis, as previously observed for regions losing TF binding during division^**10**,**11**^. Therefore, the nucleosome organisation capacity of ESRRB is also shared by NR5A2.

**Fig. 4.**
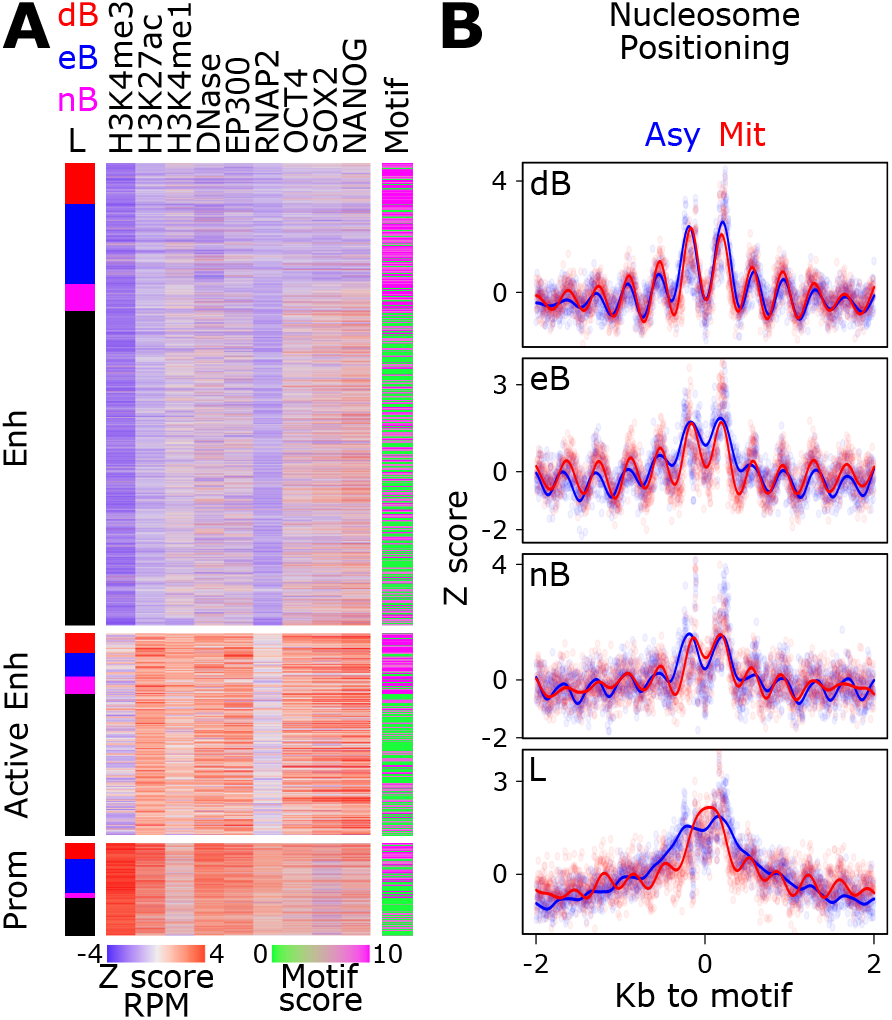
The chromatin status of ESRRB/NR5A2 binding regions. **(A)** Enrichment levels of marks of activity (H3K4me3, H3K27ac, H3K4me1, DNase accessibility, EP300, RNAP2) as established by Encode and characteristic of Enhancers (Enh), Active Enhancers (Active Enh) or promoters (Prom), as well as of TF binding (OCT4, SOX2 and NANOG) across all ESRRB/NR5A2 binding regions. On the left, the attribution as dB, eB, nB and L is shown, on the right, the mean motif score per region. **(B)** Average nucleosome positioning profiles across dB, eB, nB and L regions in asynchronous (Asy) and mitotic (mit) cells. The dots represent raw data and the lines a Gaussian Process Regression.

### ESRRB/NR5A2 mark rapidly reactivated genes

We next aimed at establishing whether ESRRB/NR5A2 binding, particularly during mitosis, is associated with post-mitotic gene transcription dynamics. To simplify these analyses, we focused on three binding groups: regions book-marked by 2 factors (dB), regions bookmarked by a single factor (B) and regions losing both factors in mitosis (L). We considered a gene as a target when either a 2kb-long window centred on its transcription start site (TSS) or an enhancer to which it had been previously linked using 3D conformational data and epigenomic analyses^**36**^ overlapped with ESRRB/NR5A2 binding regions. This association was moreover hierarchical: when a gene promoter or enhancer was overlapped by a dB region it was labelled as dB and the remaining genes were subsequently labelled, in order, as B, L or unbound. This led to gene groups of similar size: dB, 3529; B, 2594; L, 2671; unbound 5163. Next, we plotted the post-mitotic transcription dynamics of these 14,000 genes **(Fig.5A)** using highly temporally resolved data^**9**^, and computed the proportion of genes in each dB/B/L/unbound category for sliding windows of 1000 genes (step=10) displaying continuously increasing reactivation intensities. We observed that ESRRB/NR5A2 binding regions were progressively enriched as genes reactivate faster and more strongly (**Fig.5B**, left panel and **Fig.S5A**). Moreover, this progressive increase of the enrichment was solely due to regions bookmarked by both ESRRB and NR5A2 (dB, **Fig.5B**, right panel). This indicates that the combined mitotic bookmarking by ESRRB and NR5A2 may drive the efficiency of post-mitotic gene transcription. To assess this more quantitatively, we calculated the mean transcription profile of dB, B and L genes and compared them to unbound genes: all 3 groups of ESRRB/NR5A2-bound genes reactivated faster and more drastically than genes not bound by the two nuclear receptors (**Fig.5C**, left panel). Moreover, dB genes were by far the most efficiently reactivated genes, followed by B and L genes, which displayed relatively similar kinetics. Splitting the associations by the type of regulatory element further showed that genes bookmarked by ESRRB and NR5A2 reactivate more robustly than unbound genes, whether they bind at promoters or at enhancers **(Fig.S5B)**. However, only promoters showed some level of increased reactivation when bookmarked by a single nuclear receptor as compared to lost or unbound promoters **(Fig.S5B)**. Altogether, these observations support the notion that bookmarking factors promote gene reactivation in daughter cells, especially when two TFs are involved. The effect of double mitotic bookmarking by ESRRB and NR5A2 was moreover observed both for genes that are and that are not targeted by other TFs such as NANOG, OCT4 and SOX2 (**Fig.5B**, middle and right panels). Having established that mitotic bookmarking by ESRRB and NR5A2 is associated with strong post-mitotic gene reactivation, we aimed at providing functional evidence of such correlation. For this, we tagged each of the alleles of *Esrrb* and *Nr5a2* with the Auxin-induced degradation domain, which we previously used to efficiently degrade CTCF in mitosis and upon release into interphase^**9**,**11**^. Accordingly, treatment with the auxin analogue 5-Ph-IAA (thereafter IAA) led to an important reduction of both ESRRB and NR5A2 protein levels **(Fig.S6A)**. Therefore, we proceeded to synchronise the cells in mitosis with a two-step approach, first inhibiting CDK1 to enrich the population in G2, and then inhibiting microtubule dynamics to arrest cells in pro-metaphase. During the second step we added IAA to initiate ESRRB/NR5A2 degradation as the cells enter mitosis and kept it throughout the release into the next interphase, as described^**9**^. We collected multiple time-points after mitosis and prepared total ribodepleted RNA-seq libraries to quantify pre-mRNAs as a proxy of transcriptional activity^**9**^ **(Table S4)**. Contrary to our expectations, we did not observe major differences in the global reactivation dynamics of ESRRB/NR5A2-depleted cells compared to their respective control **(Fig.S6B-D)**.

**Fig. 5.**
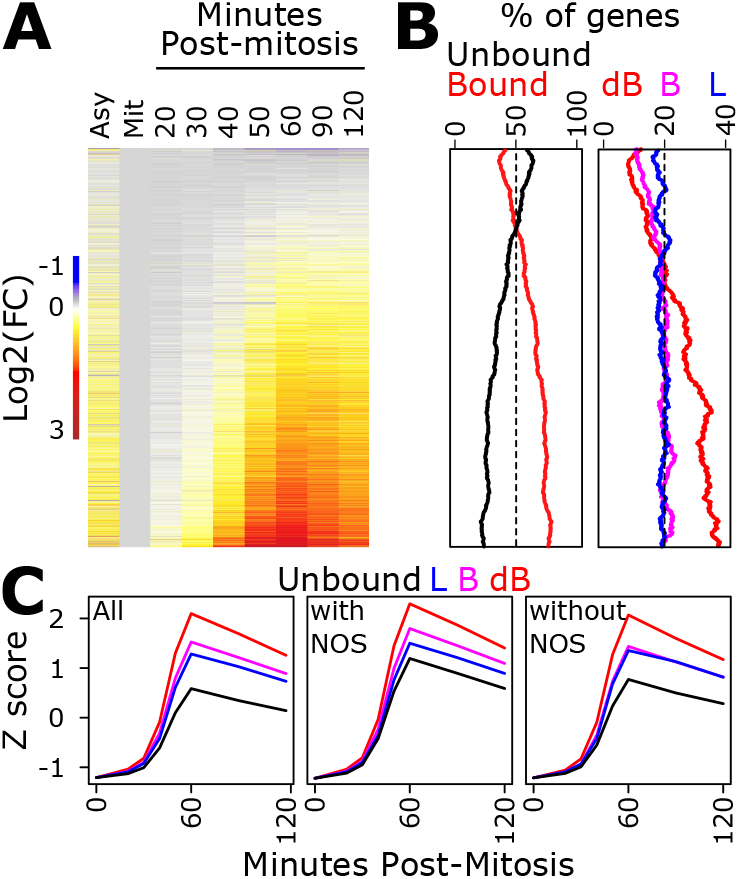
Mitotic bookmarking by ESRRB and NR5A2 is associated with fast post-mitotic gene reactivation. **(A)** Heatmap of post-mitotic transcription levels ordered by increasing mean reactivation. **(B)** Percentage of genes bound by ES-RRB/NR5A2 (left) and assigned to dB, B and L regions (right), for groups of 1000 genes sliding down the heatmap, with a step of 10 genes. **(C)** Average post-mitotic reactivation dynamics of dB, B or L. On the left, all genes, on the middle and the right those also bound, or not, by NANOG or OCT4 or SOX2 (NOS).

### ESRRB and NR5A2 activate genes associated with pluripotency during the M-G1 transition

While the global effects of the double depletion of ESRRB/NR5A2 are minor, further exploration of the dataset enabled us to identify two PCA dimensions (PC5 and PC6) describing clear differences between IAA-treated and control cells, even if capturing a small proportion of the total variance **(Fig.6A)**. Extraction of the genes contributing mostly to PC5 and PC6 identified two groups that were either down or upregulated upon IAA treatment **(Fig.6B)** and displayed concordant regulation in independent datasets generated in inducible ESRRB/NR5A2 double-KO ES cells^**14**^ **(Fig.S6E)**, indicating that they represent bona-fide ESRRB/NR5A2-responsive genes. Notably, while downregulated genes responded throughout the whole time-course analysis, upregulated genes only responded at the latest stages, suggesting that the primary effect of ESRRB/NR5A2 during the M-G1 transition is to activate transcription. Gene ontology and gene set enrichment analyses showed that downregulated genes were strongly enriched in members of the pluripotency network, and to a lesser extent in metabolic pathways **(Fig.6C)**. Indeed, several pluripotency TFs were found down-regulated after mitosis, albeit at variable levels, with some being more affected than others by ESRRB/NR5A2 depletion **(Fig.6D)**. Prompted by these results, we aimed at comprehensively identifying ESRRB/NR5A2 responsive genes after mitosis, by using direct statistical comparisons between control and IAA treated cells, as well as by taking advantage of previously identified targets^**14**^. We found 1013 genes that were downregulated after mitosis in the absence of ESRRB/NR5A2 and 941 that were upregulated, which displayed a similar behaviour **(Fig.S6F)** and functional associations **(Table S5)** to those observed in the more restricted gene sets extracted from PCA analysis **(Fig.6B,C)**. Importantly, we also found a strong association between the ESRRB/NR5A2 binding status and the gene’s responsiveness to ESRRB/NR5A2 depletion (Chi-squared p.value=5.84e-77), with nearly 80% of downregulated genes being bound at known regulatory elements **(Fig.6E)** and around 50% being bookmarked by both ESRRB and NR5A2 **(Fig.6E)**, which represents a strong and significant association **(Fig.6F)**. In contrast, upregulated or non-responsive genes were partially depleted of double bookmarked sites **(Fig.6E,F)**. We conclude that the combined bookmarking activity of ESRRB and NR5A2 at promoters and enhancers primarily fosters the activation of around 1000 genes associated with pluripotency, including important regulators such as *Tfcp2l1, Tbx3, Nanog* or *Klf5*. Finally, we analysed the expression dynamics of ESRRB/NR5A2 responsive genes after mitosis during embryoid body differentiation^**37**^ and early mouse embryogenesis^**38**,**39**^. We found that genes activated by ESRRB/NR5A2 after mitosis are rapidly downregulated upon in-vitro differentiation (**Fig.6G** left and **Fig.S6G**) and, conversely, transiently upregulated in the naïve pluripotent compartments of the early blastocyst (**Fig.6G** right and **Fig.S6H**). Conversely, genes repressed by ESRRB/NR5A2 after mitosis are globally upregulated during differentiation (**Fig.6G** left and **Fig.S6G**) and, in-vivo, maternally inherited, cleared and then re-expressed at E5.5 (**Fig.6G** right and **Fig.S6H**), when the dismantlement of naïve pluripotency starts. These observations provide further support to the biological significance of the genes mitotically bookmarked and activated by ESRRB/NR5A2 during the M-G1 transition.

**Fig. 6.**
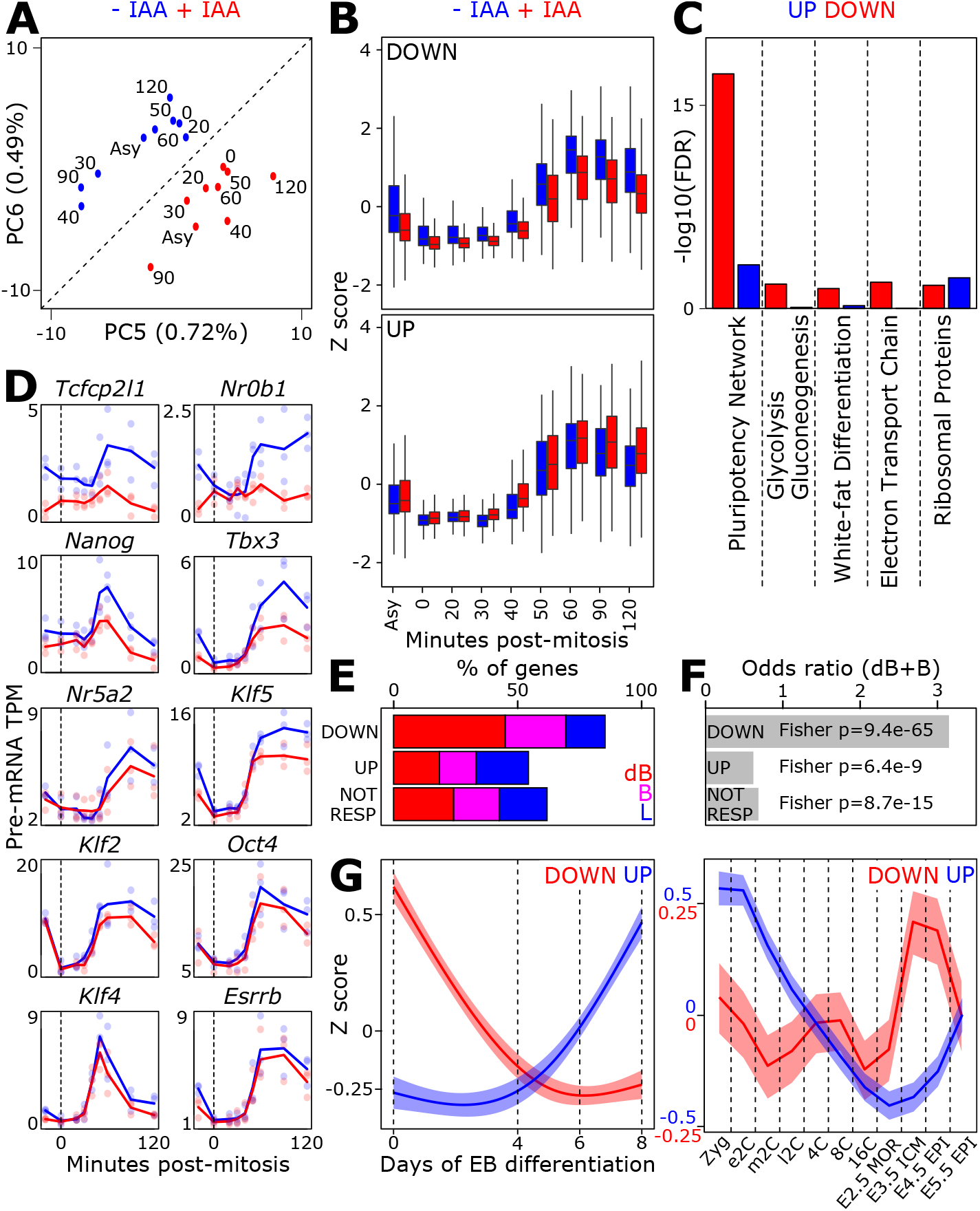
Mitotic bookmarking by ESRRB/NR5A2 promotes the reactivation of genes associated with pluripotency. **(A)** PCA analysis of post-mitotic gene transcription dynamics of cells exiting mitosis in either the presence (-IAA) or absence (+IAA) of ESRRB/NR5A2. PC5 and PC6 capture ESRRB/NR5A2-dependent differences. **(B)** Post-mitotic gene transcription dynamics of genes derived from the loadings of PC5/PC6 and displaying either a downregulation or an upregulation upon ESRRB/NR5A2 depletion. **(C)** Gene ontology analyses of the gene groups shown in (B). **(D)** Examples of post-mitotic gene reactivation dynamics for genes belonging to the pluripotency network, in the presence (blue) or the absence (red) of ESRRB/NR5A2). **(E)** Percentage of genes associated with ESRRB/NR5A2 dB, B (eB+nB) or L regions, for extended sets identified as downregulated, upregulated or not responsive. **(F)** Statistical analysis of the association of the sets shown in (E) with bookmarked regions (dB+eB+nB). **(G)** Mean expression profile during Embryoid Body differentiation (left) or Early Mouse Embryogenesis (right) of genes downregulated or upregulated upon ESRRB/NR5A2 depletion during the M-G1 transition (Zyg: Zygote; e/m/l2C: early/mid/late 2 cell stage; 4/8/16C: 4/8/16 cell stage; MOR: Morula; ICM: Inner Cell Mass; EPI: Epiblast.

## Discussion

In this study we have identified a family of gene regulators, nuclear receptors, as potentially common mitotic bookmarking factors, and have focused on two, ESRRB and NR5A2, to further characterise their binding throughout mitotic chromatin and their function in post-mitotic gene reactivation. While these factors are structurally related and their binding profiles highly similar, they can display different biophysical behaviours. These two TFs also display subtle differences in terms of the DNA binding motifs with which they preferentially interact, which become particularly relevant in the context of mitosis where the chromatin poses additional constraints for TF binding as compared to interphase. Notwithstanding, they functionally work together to maintain ordered nucleosomal arrays in mitosis and promote gene reactivation in daughter cells. Although globally their impact is not drastic, indicating that additional mechanisms may contribute to the fast reawakening of the genome, the two factors specifically promote transcription of a large subset of their mitotic targets during the M-G1 transition. Crucially, these responsive genes are strongly associated with fundamental regulatory processes supporting cell identity.

The pluripotency network is composed of a plethora of TFs that promote self-renewal and preserve pluripotency. This network is believed to be robust to the loss of single factors, since only the individual depletion of OCT4 or SOX2 leads to drastic differentiation^**40**,**41**^. This is in part due to TF redundancy, as computationally predicted^**42**^ and as shown for KLF2/KLF4/KLF5^**43**^ or for ESRRB/NR5A2^**14**^. Nevertheless, self-renewal necessarily involves undergoing mitosis, which represents an obstacle to the regulation mediated by most TFs^**1**^. Therefore, specific mechanisms may have evolved to ensure the regulatory continuity throughout cell generations and to prevent inappropriate escape from pluripotency. Mitotic bookmarking by ESRRB and NR5A2 offers, in this regard, a suitable mechanism to facilitate the reassembly of functional regulatory complexes after mitosis and promote target gene reactivation. This scenario is supported by the fact that both ESRRB and NR5A2 are simultaneously required to properly recruit NANOG, OCT4 and SOX2 at a large number of regulatory elements, notably enhancers^**14**^, and by the identification of these factors, among others, in the chromatin-associated and ESRRB-centred proteome in interphase. Nevertheless, the consequences of the loss of ESRRB/NR5A2 during the M-G1 transition are, for most genes, relatively modest, except for prominent examples such as *Tfcp2l1, Tbx3* or *Nanog*. This may be ascribed to several reasons. First, the global burst of transcription taking place after mitosis^**44**^, which is particularly strong in ES cells^**9**^, may be robust enough to be insensitive to the loss of ESRRB/NR5A2. Second, our analyses may still be compromised by additional bookmarking factors cooperating with ESRRB/NR5A2. Indeed, it is noteworthy that both the short motif identified at regions exclusively bookmarked by ESRRB, as well as the long motif identified at NR5A2-bookmarked regions, comprise the core nuclear receptor binding consensus and in particular match the ESRRA and RXRB motifs reported by the Jaspar database, respectively **(Fig.S3C)**. Since both ESRRA and RXRB are present in the mitotic ESRRB-proteome reported here, additional redundancy among nuclear receptors may further attenuate the correlations and the level of changes observed upon ESRRB/NR5A2 depletion. Thus, it might be argued that post-mitotic transcription is driven both by general and yet to be characterised mechanisms responsible for the global transcriptional burst, but also by cohorts of redundant mitotic bookmarking activities that impart specificity to the kinetics and amplitude of gene reactivation. Notably, the genes more dramatically influenced by ESRRB/NR5A2 mitotic book-marking include actors of particular importance for ES cell identity, as also suggested by the specificity in their developmental pattern of expression. This might be a general characteristic of the genes responding during M/G1 to the action of cell type-specific mitotic bookmarking TFs, as supported by the recurrent identification of cell identity genes within repertoires of mitotic bookmarking targets^**3-5**^.

Redundancy by nuclear receptors may also be the reason why genes losing ESRRB/NR5A2 binding in mitosis still reactivate more efficiently than those that are not bound at all. Alternatively, the inheritance of the fraction of ESRRB/NR5A2 that is not chromatin-bound during division may also be important to activate target genes rapidly in G1. In accord with this view, the binding by TFs that do not act as mitotic bookmarking factors in our cells, such as OCT4, SOX2 and NANOG, is also associated with accelerated transcription patterns after mitosis in comparison with unbound genes. In this regard, the differential contribution of promoter or enhancer-bound TFs in interphase and mitosis appears relevant. Genes exclusively bound either at the promoter or at cognate enhancers by ESRRB/NR5A2 display distinct properties: those losing promoter binding by ESRRB/NR5A2 in mitosis do not reactivate faster than unbound genes, whereas genes losing enhancer binding do. This suggests that promoters may be subject to additional mechanisms fuelling their post-mitotic activation, such as particularly well-preserved accessibility^**45**^, mitotic bookmarking by TBP^**8**^ or by CTCF^**9**^, or APC/C-driven regulations^**46**^. Hence, in this permissive context the additional contribution of ESRRB/NR5A2 might have a minor impact. In contrast, enhancer activity, which ultimately depends on the assembly of enhanceosomes constituted by different TFs, may display higher sensitivity to the order and the kinetics of rebinding. Hence, given the role of ESRRB/NR5A2 in enabling enhancer occupancy by other pluripotency TFs in interphase^**14**^ and the pioneering activity shown for NR5A2^**47**^, it is likely that both their mitotic bookmarking activity as well as their prompt binding after mitosis may promote, together with other nuclear receptors, the perpetuation of a chromatin configuration that facilitates the fast reactivation of most pluripotency-associated genes. Additional studies are now required to dissect the underlying mechanisms, which likely involve the capacity of ESRRB/NR5A2 to order nucleosomes or to retain specific members of the protein complexes with which they cooperate to control transcription in interphase, identified here by proteomics. In this regard, the potential association of BRG1 (SMARCA4) and CHD4, the catalytic subunits of two nucleosome remodelling complexes important for ES cells^**48**,**49**^ may contribute to both the maintenance of nucleosome organisation and to promote the reassembly of full regulatory complexes in interphase via protein-protein interactions. Finally, besides redundantly cooperating to reinstate enhancer and promoter activity, ESRRB, NR5A2 and possibly other nuclear receptors, might directly take part in fundamental mitotic processes, as suggested by the association on mitotic chromatin of ESRRB with proteins involved in the regulation of mitosis. In this sense, nuclear receptors might emerge as holistic regulators of the ability of cells to progress through cell division.

The observation that multiple nuclear receptors and associated factors are likely engaged in mitotic bookmarking of key regulatory elements of the ES cell genome suggests that this super-family of regulators may have a constitutive role in post-mitotic gene reactivation in other cell types. Indeed, several nuclear receptors have been suggested to act as mitotic bookmarking factors, mainly based on their capacity to coat mitotic chromosomes^**19-25**^, including the Androgen Receptor, the Estrogen Receptor Alpha, the Pregnane X Receptor, the Vitamin D Receptor or the Hepatocyte Nuclear Factor 1 Beta. Given the importance of these regulators in cell homeostasis and in several pathological conditions, we believe that globally assessing the mitotic bookmarking role of the entire family of nuclear receptor requires new attention. Showcasing the potential clinical implications, it has been proposed that mutations associated with pathological outcomes specifically alter the chromosomal retention of these proteins^**22**,**50**^. Despite this, a key question remains: what is the advantage of having a large battery of nuclear receptors acting as mitotic bookmarking factors? Since most of these receptors are activated in a ligand-dependent manner, it is likely that mitotic bookmarking may provide cells with a conditional memory of gene activity that is directly connected to the cellular environment, in a manner much more flexible and amenable to rapid regulatory changes than more canonical means of epigenetic inheritance. Of course, ESRRB and NR5A2 are orphan nuclear receptors, which do not depend on specific ligands given that their LBDs constitutively reside in an active configuration^**51-53**^. Yet, it is possible that the ancestral proteins from which they derived during evolution had an environmental-sensing function^**54**^, including for their mitotic bookmarking activity as supported by the agonist-dependent mitotic retention of the Androgen Receptor^**20**^. Losing the dependency to a ligand, while conserving a mitotic bookmarking activity, may therefore be linked to the co-option of orphan receptors as constitutive regulators and developmental determinants^**55**^. Studying the mitotic bookmarking capacity of nuclear receptors, in light of their evolutionary history, may thus inform on how the establishment of mechanisms of gene expression memory transit from being driven by the environment to become constitutively operational in specific cell types or developmental stages.

## Methods

### Cell culture and mitotic preparations

ES cells – E14Tg2a, EKOiE^**14**^, EKOie-NrKO^**14**^, FLAG-Nr5a2^**14**^, NR5A2-GFP/ESRRB-mCherry^**14**^, ESRRB/NR5A2-GFP lines and ESRRB/NR5A2-IAA ESCs (see details below) – were cultured on serum and LIF conditions as previously described^**11**^. Mitotic ES cells (>95% purity as assessed by DAPI staining and microscopy), were obtained using a double synchronisation method based on the CDK1 inhibitor RO-3306 (10 μM; Sigma, SML0569), nocodazole (50ng/ml; Sigma, M1404) and shake-off, as previously described^**9**^. For post-mitosis analyses, cells were seeded in separate dishes (one per time-point), purposely uncoated with gelatine, and lysed in cold TRIzol (ThermoFisher, 15596026) 20, 30, 40, 50, 60, 90 and 120 min after release from the mitotic block^**9**^. ESRRB/NR5A2 depletion was achieved with 0.5 mM auxin (5-Ph-IAA BioAcademia, 30-003), added during the 5h of nocodazole block and maintained during the whole post-mitotic release. Asynchronous cells were treated in parallel during 5h.

### ES cell derivation

ESRRB/NR5A2-GFP cells were generated by stable transfection of a CAG-driven vector expressing C-terminal fusions of ESRRB or NR5A2 variants to GFP (connected by a Glycine linker and linked to an IRES-Puromycin resistance cassette) and selection of single clones. NR5A2 variants included the full protein from ENSEMBL transcript *Nr5a2-205* (Uniprot Q1WLP7) and two truncated version coding for the DNA binding domain alone or for a fragment spanning the DNA binding domain and all the remaining C-terminal portion of the protein (aa: DEDLEE … LHAKRA). For experiments shown in Fig.2B, similar expression constructs were transiently transfected by lipofection in EKOiE, or EKOie-NrKO cells. ESRRB/NR5A2-AID cells were generated by CRISPR/Cas9, first inserting a CAG-OsTir2-T2a-NeomycinR cassette at the TIGRE locus (gRNA 3-ACTGCCATAACACCTAACTT-5), then a LoxP-PuromycinR-LoxP-HA-AID-Gly5 cassette at the start codon of *Nr5a2* (ENSEMBL transcript *Nr5a2-205*; gRNA 5-CCACTTTGGGCAGCATGACA -3), and finally a LoxP-PuromycinR-LoxP-3xFLAG-Gly5 cassette at the start codon of *Esrrb* (ENSEMBL transcript *Esrrb-206*; gRNA 5-TGAACCGAATGTCGTCCGAC-3). After each round, single colonies were expanded and cells homozygous for correctly targeted alleles identified by PCR on genomic DNA and sequencing. In addition, after each insertion of the AID degron the selection cassette was removed by Cre-mediated recombination.

### Proteomics

Asynchronous and mitotic cells were obtained during successive experimental rounds, fixed with DSG (2mM; Sigma, 80424-5mg) for 50min at RT followed by 10min with formaldehyde (1%, Thermo, 28908), sonicated as previously described^**10**^ and stored at -80°C until 300.10^6^ cells for each were accumulated. Next, ESRRB and control immunoprecipitations (IP) were performed in parallel in triplicates using 50.10^6^ cells per IP and a standard ChIP procedure with anti-ESRRB (Perseus Proteomics, H6-705-00) and control antibodies, except that after the last wash the beads were resuspended in 2x LDS buffer/100mM DTT, incubated for 35 min at 95°C while shaking and spun for 10 min at RT at maximum speed. Samples were stored at - 20°C until further processing. The eluates, after equilibrating their temperature to RT, were alkylated by incubating with 5.5 mM chloroacetamide for 30 min in the dark and then loaded onto 4-12% gradient SDS-PAGE gels. Proteins were stained using the Colloidal Blue Staining Kit (Life Technologies). Due to DSG/FA crosslinking, the proteins appeared as a smear upon SDS-PAGE electrophoresis; therefore, the bands corresponding to heavy and light chains of the IP antibodies were not cut out to avoid losing potentially relevant proteins. All proteins were digested in-gel using trypsin.

Peptides were extracted from gel and desalted on reversed phase C18 StageTips. Peptide fractions were analyzed on a quadrupole Orbitrap mass spectrometer (Q Exactive Plus, Thermo Scientific) equipped with a UHPLC system (EASY-nLC 1000, Thermo Scientific). Peptide samples were loaded onto C18 reversed phase columns (15 cm length, 75 μm inner diameter, 1.9 μm bead size) and eluted with a linear gradient from 8 to 40% acetonitrile containing 0.1% formic acid in 2-hours. The mass spectrometer was operated in data dependent mode, automatically switching between MS and MS2 acquisition. Survey full scan MS spectra (m/z 300 – 1700) were acquired in the Orbitrap. The 10 most intense ions were sequentially isolated and fragmented by higher-energy C-trap dissociation (HCD). An ion selection threshold of 5,000 was used. Peptides with unassigned charge states, as well as with charge states less than +2 were excluded from fragmentation. Fragment spectra were acquired in the Orbitrap mass analyser. Raw data files were analysed using MaxQuant (development version 1.5.2.8)^**56**^. Parent ion and MS2 spectra were searched against a database containing all mouse protein sequences obtained from the UniProtKB released in 2016 using Andromeda search engine^**57**^. Spectra were searched with a mass tolerance of 6 ppm in MS mode, 20 ppm in HCD MS2 mode, strict trypsin specificity and allowing up to 3 miscleavages. Cysteine carbamidomethylation was searched as a fixed modification, whereas protein N-terminal acetylation and methionine oxidation were searched as variable modifications. The dataset was filtered based on posterior error probability (PEP) to arrive at a false discovery rate of below 1% estimated using a target-decoy approach^**58**^. Table S1 reports all the proteins identified by MS in triplicate control and ESRRB IPs in asynchronous and mitotic cells, along with quantifications, normalized intensities and additional metrics. Razor and unique peptides were used to compute a fold enrichment and p-value between the ESRRB and control IP in either asynchronous or mitotic cells using the DEseq package^**59**^. Proteins displaying a fold change above 5 and a p-value below 0.05 in either asynchronous or mitotic cells were selected for further analyses. This list was filtered based on ‘Reverse’ hits, ‘Contaminants’ and ‘identified by site’ parameters. All detected immunoglobulins as well as other proteins belonging to the top 1000 frequently identified proteins in MS datasets (https://www.thegpm.org/lists/index.html) were ignored. This led us to 105 proteins identified as associated with ESRRB in either asynchronous or mitotic cells, available in Fig.S1 and in Table S2. Proteins displaying a p-value below 0.05 were considered as positive mitotic hits. However, we further classified them as high or low confidence depending on their general abundance in mitotic cells and their identification after Formaldehyde-only fixation (FA). To do this, we performed an additional round of MS using the mitotic replicate 2 for ESRRB IP together with its corresponding input (DSG/FA fixed) as well as an IP/Input generated in parallel after FA fixation. Proteins displaying higher Razor and unique peptides in both the DSG/FA and FA IPs compared to their respective inputs were considered of high confidence (Table S2). These 105 hits were analysed using the String database^**60**^: first, all proteins were used as an input to identify functionally related groups (subnetworks) based on Ontology annotations; second, all possible functional and biochemical interactions between each protein and the rest was computed; third, subnetworks were connected using the sum of all the interaction scores existing between all individual proteins of each group.

### Imaging

For immunofluorescence, cells were plated on IBIDI hitreat plates coated overnight with poly-L-ornithine 0.01% (Sigma, Cat P4957) at 4°C, washed and coated 2h with 10μg/ml laminin (Millipore, Cat CC095). Fixation and immunofluorescence was performed with either PFA or DSG+PFA, as described^**10**^, using 2 μg/ml mouse monoclonal anti-flag for ESRRB (M2 Sigma, F3165) and 1μg/ml polyclonal rabbit anti-HA for NR5A2 (Abcam, ab9110) antibodies. Images were acquired with a LSM900 Zeiss microscope using a 64× oil-immersion objective. For live imaging and FRAP analyses, ES cells expressing fluorescent protein fusions were grown on IBIDI plates, incubated with 250 nM Hoechst-33342 for 30 min before imaging and imaged at 37°C in a humidified atmosphere (7% CO2). Images were acquired with a 63× oil immersion objective on a Nikon Ti2E equipped with a Yokagawa CSU W1 spinning disk module and a Photometrics sCMOS Prime 95B camera. For FRAP, fluorescence recovery was analysed every 0.3-1s (NR5A2) and 50-100 ms (ESRRB) after photobleaching in Matlab as described previously^**61**,**10**^ and the plots corrected to min=0 and max=1.

### Identification of ESRRB/NR5A2 binding sites displaying distinct behaviors in mitosis

NR5A2 ChIP-seq was performed using mitotic cells as previously described and in parallel with already published datasets generated in asynchronous cells^**14**^. Briefly, cells were crosslinked with DSG (2mM; Sigma, 80424-5mg) for 50min at RT followed by 10min with formaldehyde (1%, Thermo, 28908). Cells were then sonicated with a Bioruptor Pico (Diagenode) and immunoprecipitated with anti-flag antibodies (M2 Sigma, F3165). Precipitated DNA was used for library preparations^**10**^ and sequenced externally by Novogene Co Ltd. Reads were aligned with Bowtie2^**62**^ to the mm10 genome; only those with a single discovered alignment were kept. Peaks were called against relevant inputs for all mitotic samples using MACS2^**63**^ and filtered to have (1) MACS2 FDR < 0.05 in all three replicates, (2) mean enrichment over the input > 2, (3) FDR of the enrichment over the input < 0.05, calculated with a previously described generalised linear model^**10**^. The resulting mitotic NR5A2 peaks were combined with previous collections of confident ESRRB/NR5A2 peaks^**10**,**14**^ and mitotic bookmarking calls for ESRRB^**10**^. Finally, peaks were annotated as dB/eB/nB/L. The compendium of ESRRB/NR5A2 regions, their quantifications and mitotic book-marking status are available as Table S3.

### Identification of differentially expressed genes during the M-G1 transition

RNA was extracted with 500 μl TRIzol (ThermoFisher, 15596026), treated with DNAse I (Qiagen) and used for the generation of Ribo-depleted, stranded and paired-end RNA-seq libraries, prepared and sequenced by Novogene Co Ltd. Pre-mRNA levels were quantified exactly as previously described^**9**^ and those with more than 0.1 transcripts per million in at least one sample were kept for further analyses. Differentially expressed genes were obtained from two separate analyses. First, using previous lists of ESRRB/NR5A2-responsive genes^**14**^: we selected genes with an FDR < 0.05 upon the double knock-out of the two TFs, quantified their pre-mRNA levels across the datasets generated here, and identified those displaying concordant changes in plus/minus IAA during the M-G1 transition using k-means clustering. Second, using a direct comparative strategy of each time-point analysed here and keeping those with an FDR < 0.1 (DEseq2^**64**^). A fully annotated table with all genes considered, quantifications and differential expression parameters is provided in Table S4.

### Bioinformatic analyses

Analyses were performed in R (version 3.6.3). ChIP-seq quantifications were performed with the bamsignals package with systematic correction to the library sizes and counting the number of reads either falling into peak coordinates (for boxplots) or covering each base of a 2kb window centred on the middle of the peak (for enrichment heatmaps and metaplots). Boxplots and metaplots were visualized with ggplot2 package^**65**^ and heatmaps with ComplexHeatmap package^**66**^. To characterise the regions as promoters (high H3K4me3), active enhancers (high H3K27ac) or enhancers (high H3K4me1), available ES cell data from the Encode consortium was used. All additional histone modifications and DNase accessibility datasets were downloaded from Encode. ATAC-seq as well as ChIP-seq for NANOG, OCT4 and SOX2 were previously published^**10**,**67**^. All quantifications are available in Table S3. To identify the most prevalent DNA motif associated with distinct ESRRB/NR5A2 binding regions, as well as their precise genomic coordinates, we used RSAT^**68**^ with the command - markov auto -disco oligos -nmotifs 1 -minol 6 -maxol 8 - merge_lengths -2str -origin start -scan_markov 1. For each motif, the presence of a T, a C or an A/G at the 7th position of the Estrogen Related Receptor consensus was manually annotated. The information relative to the motifs can be found in Table S3. Nucleosome positioning plots were generated considering midpoints of 140-200bp fragments from published MNase-seq paired-end datasets^**10**^, after correcting the MNase-driven bias with a kmer approach and smoothing the average profiles with a Gaussian Process Regression, as described^**10**^. When multiple motifs were available per region, only one was selected to centre nucleosome positioning plots on its 5’ end; the following prioritized criteria were used: (1) best similarity score to the consensus, (2) closest distance to the middle of the peak, (3) highest enrichment for small MNase fragments (footprints <100bp). Genes were associated to the different groups of ESRRB/NR5A2 binding regions when either their promoter regions – defined as 2kb-long regions centred on the 5’ ends of known mRNA isoforms) –, or putative enhancers – defined by their epigenomic status and 3D interaction contacts as independently reported^**36**^ – overlapped with our set of ChIP-seq peaks. The approach was hierarchical, favouring dB associations over eB/nB and then L regions; genes lacking ESRRB/NR5A2 binding at these regions were qualified as unbound (Table S4). The resulting groups of genes were used to plot average reactivation trends and the proportion of dB/eB/nB/L genes displaying increased reactivation dynamics. Principal component analysis of the RNA-seq was run with the prcomp function with centred data corresponding to log2(TPM) to capture direct differences between IAA treatments. The genes more prominently contributing to PC5/PC6 were identified with the boxplot.stats function applied to the ‘rotation’ output of prcomp. Gene ontology and gene set enrichment analyses were performed using the enrichR package^**69**^, the full set of statistically significant terms can be found in Table S5. The relationships between ESRRB/NR5A2 binding at promoters/enhancers and gene responsiveness to ESRRB/NR5A2 depletion was assessed with a contingent table and Chi-squared test followed by individual two-sided Fisher’s exact tests for specific combinations of gene book-marking and responsiveness.

## Acknowledgements

The authors acknowledge Michel Cohen-Tannoudji, Thomas Gregor, Germano Cecere, Ken McElreavy and Shahragim Tajbakhsh for critical reading of the manuscript. The authors also acknowledge the Biomics, Flow Cytometry, Photonic BioImaging and Image Analysis Hub platforms of Institut Pasteur. PN acknowledges the Institut Pasteur, the CNRS, and Revive (Investissement d’Avenir; ANR-10-LABX-73) for recurrent funding and the Agence Nationale de la Recherche (ANR 20CE12002801 CHRODYNE), Ligue contre le Cancer (LNCC EL2018 NAVARRO) and the European Research Council (ERC-CoG-2017 BIND) for financial support.

## Author contributions

AC performed bioinformatic analyses with help from DF. AM performed most of the experiments with the exception of ChIP-seq and the derivation of cell lines (NF). HIB and PB performed proteomics with samples gathered by AM with help from EH. TP established the initial conditions for ESRRB IP for proteomics. FM provided help to analyse FRAP data. AD and NG provided technical help. NF and PN conceived the project and analysed the data. PN wrote the manuscript.

## Supplementary information

Six supplementary figures accompany this manuscript, they can be found at the end of this document. Four Supplementary Tables are available online.

---

**Supplementary Information, Fig.S 1.**
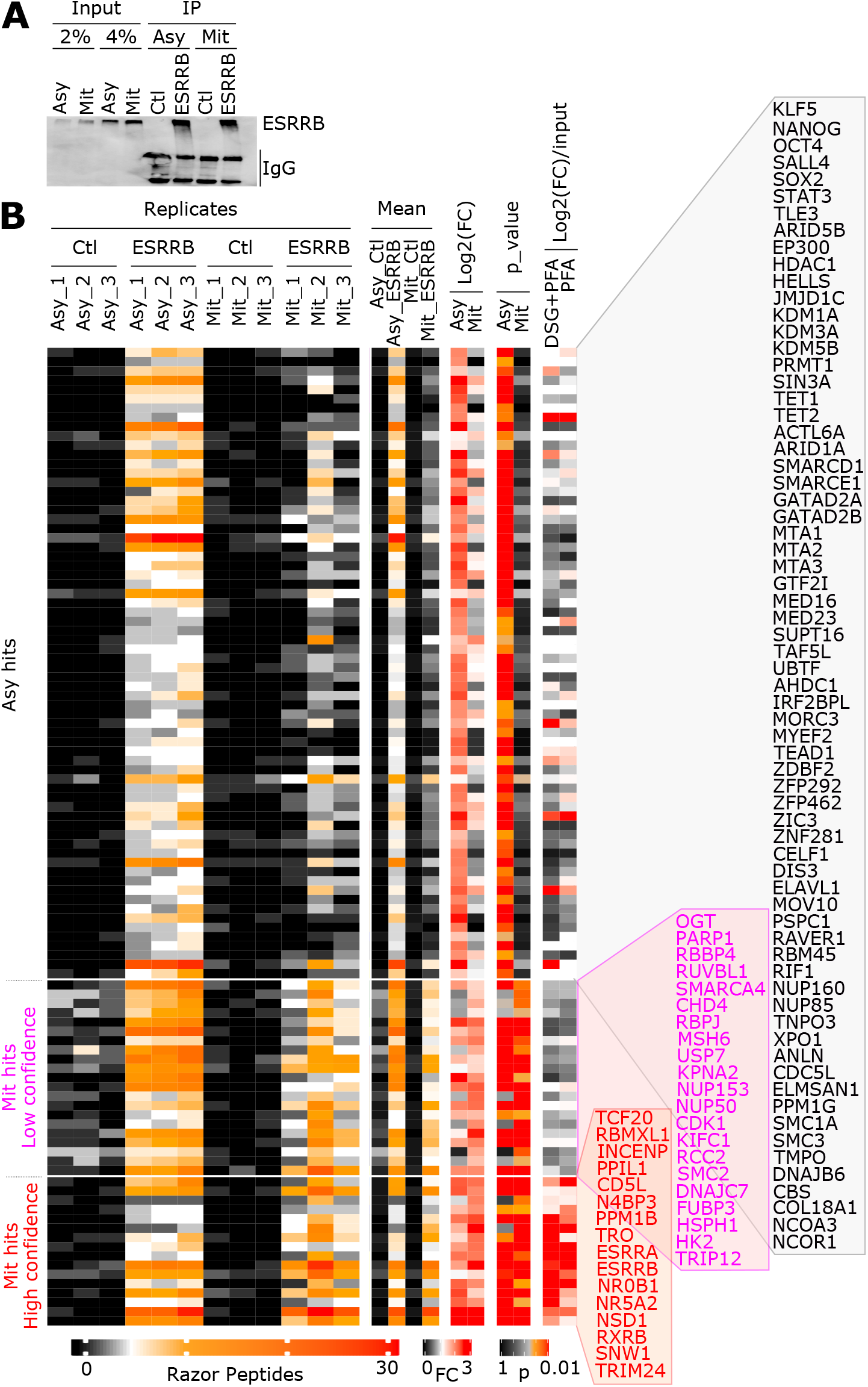
Quantifications of all protein hits in interphase and in mitosis. (A) Representative immunoprecipitation (IP) of ESRRB from DSG/FA-fixed chromatin. Asy: asynchronous cells; Mit: mitotic cells; Ctl: blank immunoprecipitation. (B) Quantifications (Razor peptides), fold changes and associated p-values used to categorise the proteomic hits as exclusive of interphase (ASY hits) or also found in mitosis (Mit hits). Two categories of mitotic hits were made, based on the fold-change of ESRRB IP versus the INPUT in mitosis (last parameters on the right of the heatmap). All the proteins belonging to each category are shown.

**Supplementary Information, Fig.S 2.**
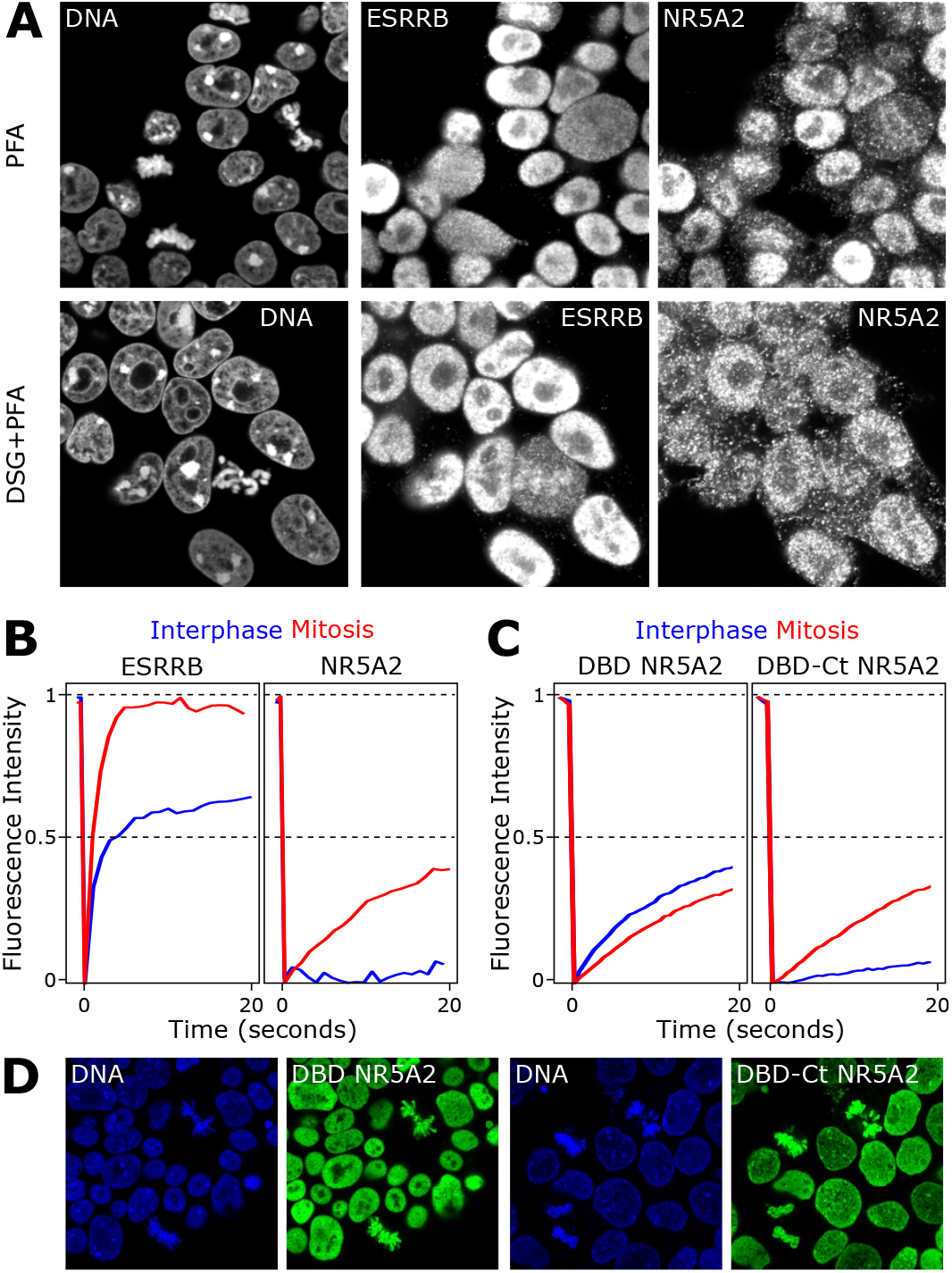
NR5A2 long-lived chromatin interactions are driven by its DNA binding domain. (A) Ilustrative example of ESRRB/NR5A2 immunostaining upon formaldehyde or DSG and formaldehyde crosslinking, as indicated. (B) Quantification of comparative FRAP assays of ESRRB-GFP and NR5A2-GFP. (C) Quantification of comparative FRAP assays of GFP fusions with the DNA binding domain of NR5A2 alone (DBD NR5A2) or with its C-terminal moiety containing the ligand binding domain (DBC-Ct). (D) Representative live imaging of the DBD or the DBD-Ct fused to GFP.

**Supplementary Information, Fig.S 3.**
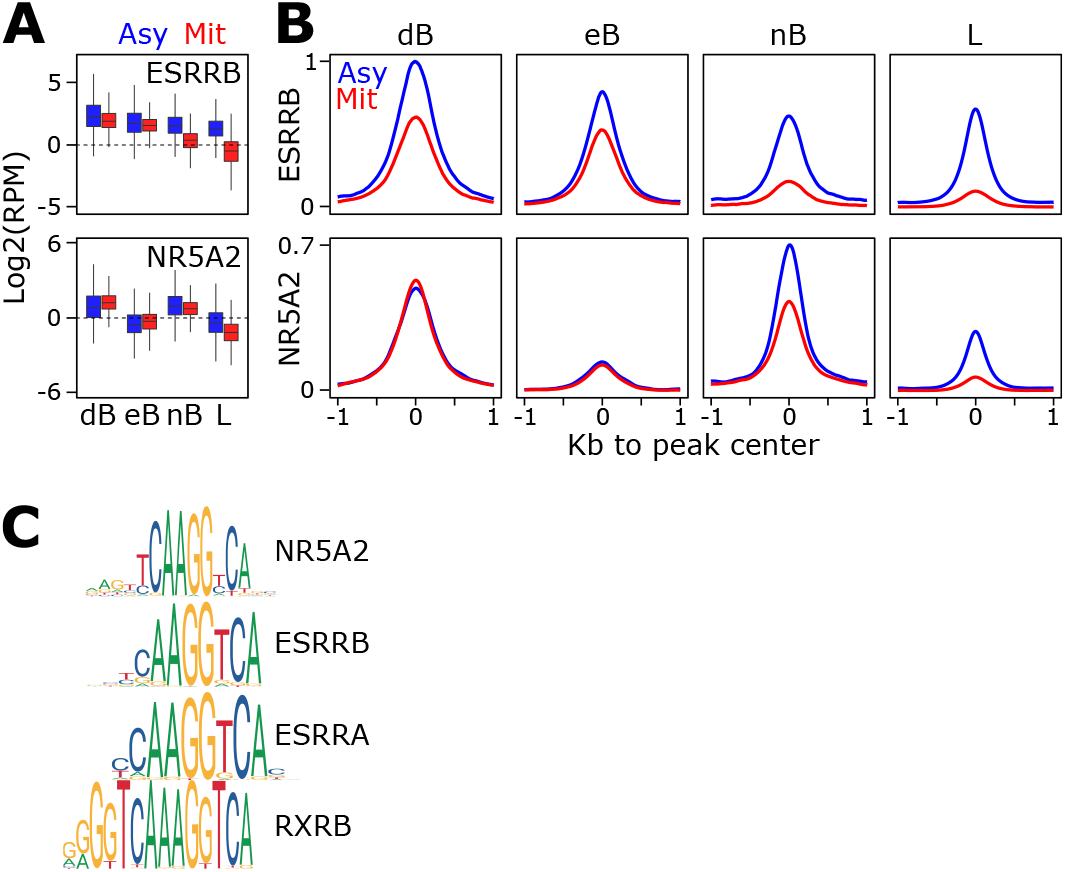
Comparison of ESRRB/NR5A2 binding levels and motifs matching the identified consensus. (A-B) Enrichment levels (Reads per million) of ESRRB and NR5A2 across the four identified clusters (dB, eB, nB, L), represented as boxplots (A) or metaplots (B). (C) Known nuclear receptors motifs (Jaspar database) matching the consensus sequence identified in the 4 binding clusters.

**Supplementary Information, Fig.S 4.**
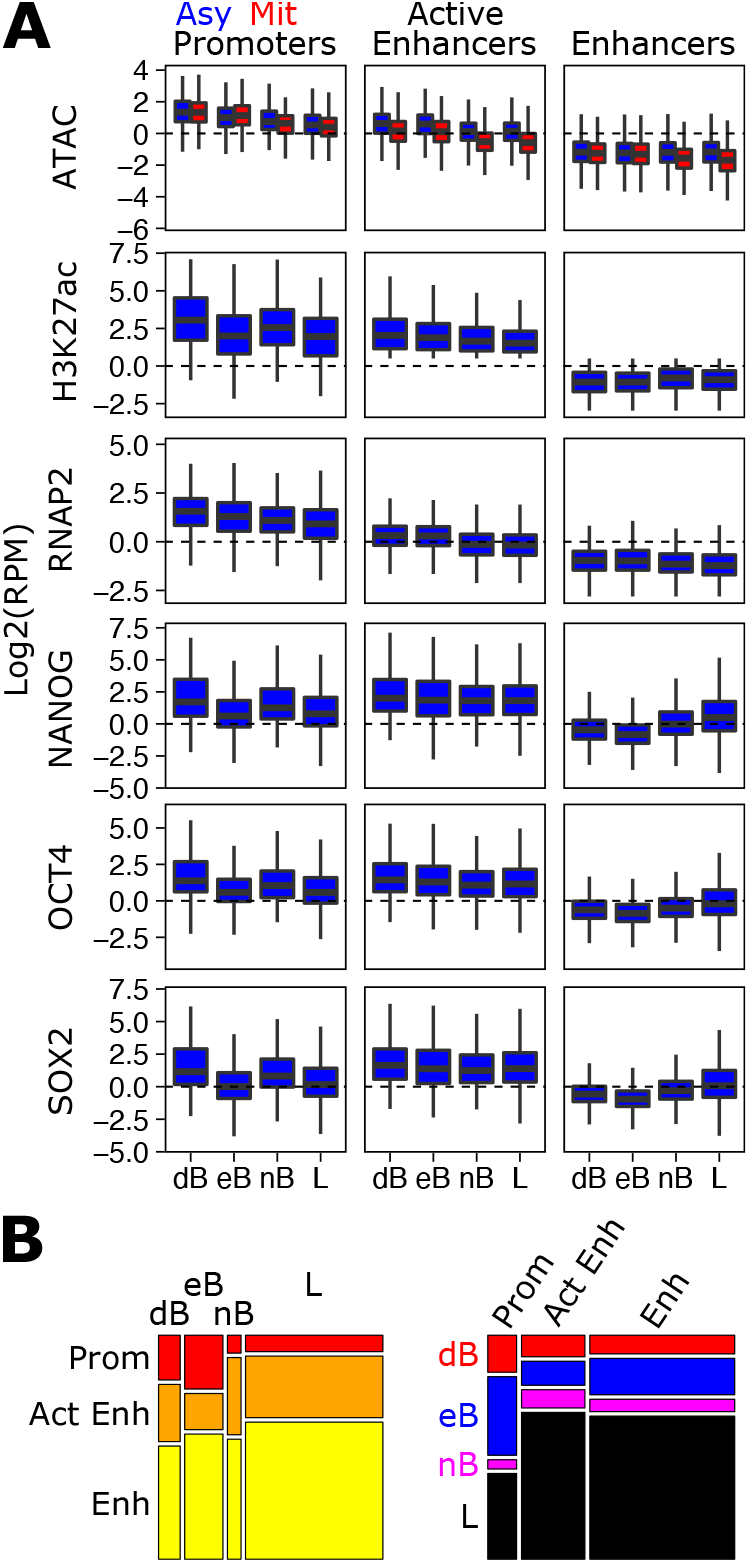
Relationships between ESRRB/NR5A2 binding and other chromatin properties at distinct gene regulatory elements. (A) Enrichment levels of the indicated factors across dB, eB, nB and L regions separated as promoters, active enhancers or enhancers. (B) Mosaic plots describing the relative proportions of promoters, active enhancers and enhancers across dB, eB, nB and L regions (left) and vice-versa (right).

**Supplementary Information, Fig.S 5.**
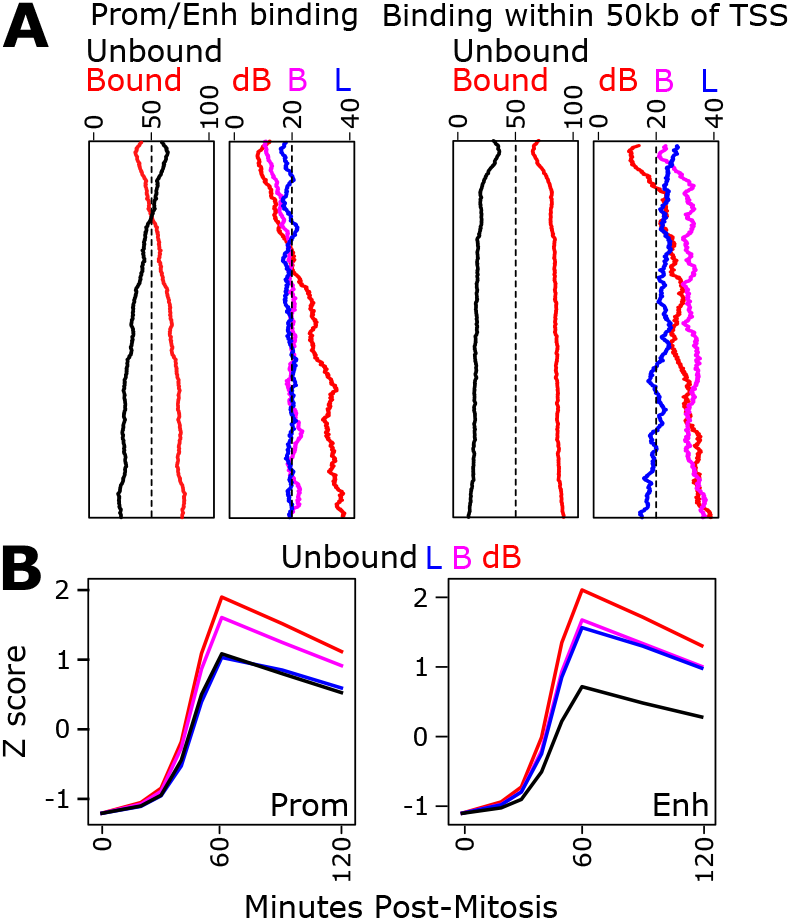
Additional correlations between ESRRB/NR5A2 and post-mitotic gene transcription. (A) Comparison of the proportion of genes identified as dB, eB, nB and L regions when the gene-region association is based on known enhancers and promoters (left, identical to Fig.5B) or when it is based on the presence of a binding regions within 50kb of the promoter (right), as previously done^**9**^. (B) Post-mitotic gene transcription dynamics when the associations consider exclusively promoters (Prom, left) or exclusively enhancers (Enh, right).

**Supplementary Information, Fig.S 6.**
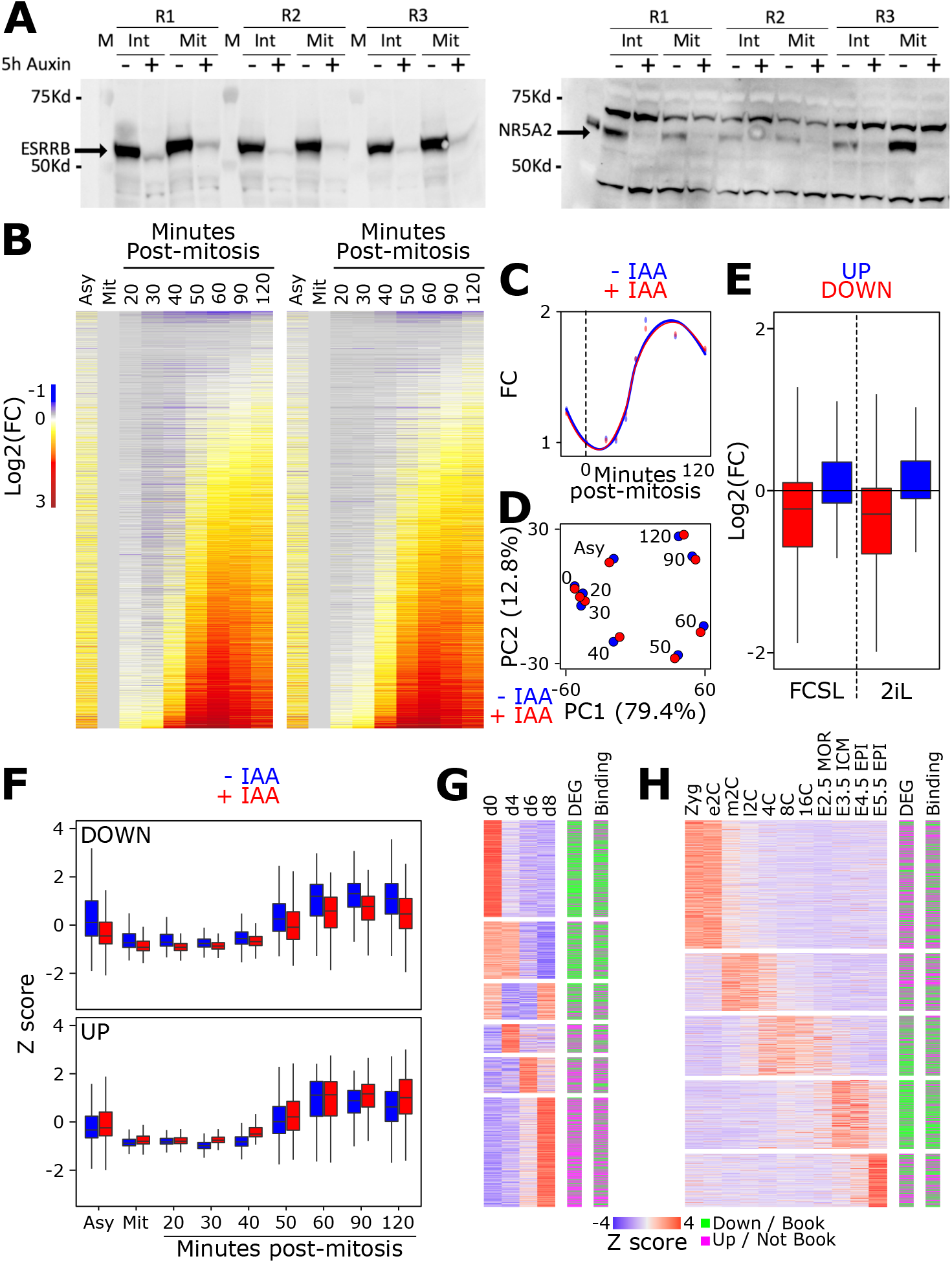
Regulation of selected genes by ESRRB/NR5A2 during the M-G1 transition. (A) Western-blot analysis of ESRRB/NR5A2 depletion upon auxin treatment in interphase and in mitosis. (B) Reactivation heatmap presented as in Fig.5A but in control (left) or ESRRB/NR5A2-depleted cells (right). (C) Global gene reactivation profile of all genes shown in (B) in presence (-IAA) or absence (+IAA) of ESRRB/NR5A2. (D) PCA analysis of post-mitotic transcription dynamics in presence (-IAA) or absence (+IAA) of ESRRB/NR5A2. (E) Fold-change of expression levels upon ESRRB/NR5A2 knock-out in two different media (FCSL or 2iL^**14**^) for the genes extracted from PC5/PC6 loadings as shown in Fig.6A, B. (F) Transcription dynamics of an extended list of genes regulated by ESRRB/NR5A2 during the M-G1 transition. (G) Expression profile of the two groups of genes shown in (F) during embryoid bodies differentiation, organised by timing of maximal expression, with the identification of the groups they belong and their mitotic bookmarking status shown on the right). (H) Identical analysis to (G) but during early mouse embryogenesis.

